# Changes of small non-coding RNAs by severe acute respiratory syndrome coronavirus 2 infection

**DOI:** 10.1101/2021.12.16.472982

**Authors:** Wenzhe Wu, Eun-Jin Choi, Binbin Wang, Ke Zhang, Awadalkareem Adam, Gengming Huang, Leo Tunkle, Philip Huang, Rohit Goru, Isabella Imirowicz, Leanne Henry, Inhan Lee, Jianli Dong, Tian Wang, Xiaoyong Bao

**Affiliations:** Department of Pediatrics, The University of Texas Medical Branch, Galveston, TX, 77555, USA; Department of Microbiology and Immunology, The University of Texas Medical Branch, Galveston, TX, 77555, USA; Department of Pathology; The University of Texas Medical Branch, Galveston, TX, 77555, USA; miRcore, Ann Arbor, MI, 48105, USA; Department of Nuclear Engineering &Radiological Sience, University of Michigan, Ann Arbor, MI, 48109, USA; Department of Computer Science, University of Michigan, Ann Arbor, MI, 48109, USA; Department of Molecular, Cellular & Developmental Biology, University of Michigan, Ann Arbor, MI, 48109, USA; The Institute for Human Infections and Immunity, The University of Texas Medical Branch, Galveston, TX, 77555, USA; The Institute of Translational Sciences, The University of Texas Medical Branch, Galveston, TX, 77555, USA

**Author notes:** Correspondence should be addressed to Xiaoyong Bao, Ph.D., Division of Clinical and Experimental Immunology & Infectious Diseases, Department of Pediatrics, 301 University Boulevard, Galveston, TX 77555-0366.

**Keywords:** SARS-CoV-2, tRF, SARS-CoV-2-derived sncRNAs

## Abstract

The ongoing pandemic of coronavirus disease 2019 (COVID-19), which results from the rapid spread of the severe acute respiratory syndrome coronavirus 2 (SARS-CoV-2), is a significant global public health threat, with molecular mechanisms underlying its pathogenesis largely unknown. Small non-coding RNAs (sncRNAs) are known to play important roles in almost all biological processes. In the context of viral infections, sncRNAs have been shown to regulate the host responses, viral replication, and host-virus interaction. Compared with other subfamilies of sncRNAs, including microRNAs (miRNAs) and Piwi-interacting RNAs (piRNAs), tRNA-derived RNA fragments (tRFs) are relatively new and emerge as a significant regulator of host-virus interactions. Using T4 PNK-RNA-seq, a modified next-generation sequencing (NGS), we recently found that nasopharyngeal swabs (NPS) samples from SARS-CoV-2 positive and negative subjects show a significant difference in sncRNA profiles. There are about 166 SARS-CoV-2-impacted sncRNAs. Among them, tRFs are the most significantly affected and almost all impacted tRFs are derived from the 5’-end of tRNAs (tRF5). Using a modified qRT-PCR, which was recently developed to specifically quantify tRF5s by isolating the tRF signals from its corresponding parent tRNA signals, we validated that tRF5s derived from tRNA GluCTC (tRF5-GluCTC), LysCTT (tRF5-LysCTT), ValCAC (tRF5-ValCAC), CysGCA (tRF5-CysGCA) and GlnCTG (tRF5-GlnCTG) are enhanced in NPS samples of SARS-CoV2 patients and SARS-CoV2-infected airway epithelial cells. In addition to host-derived ncRNAs, we also identified several sncRNAs derived from the virus (svRNAs), among which a svRNA derived from CoV2 genomic site 346 to 382 (sv-CoV2-346) has the highest expression. The induction of both tRFs and sv-CoV2-346 has not been reported previously, as the lack of the 3’-OH ends of these sncRNAs prevents them to be detected by routine NGS. In summary, our studies demonstrated the involvement of tRFs in COVID-19 and revealed new CoV2 svRNAs.

## Introduction

Severe acute respiratory syndrome coronavirus 2 (SARS-CoV-2) is a beta coronavirus belonging to the sarbecovirus subgenus of Coronaviridae family [1]. It is a positive-sense single-stranded RNA virus with a genome length of ∼30kb. Until early November 2021, the ongoing coronavirus disease 2019 (COVID-19) pandemic, caused by SARS-CoV-2, has respectively caused more than 251 million infectious cases and over five million deaths globally [2]. Symptoms after SARS-CoV-2 infection vary widely. The majority of patients are asymptomatic or mildly symptomatic, while some patients, particularly the elderly and those with other serious health problems, such as comprised immunity, diabetes, and cardiovascular/pulmonary diseases, are at the risk of developing severe symptoms and have a higher fatality rate [3]. SARS-CoV-2 is also becoming a significant health threat to the pediatric population [4]. Irrespective of disease symptoms, SARS-CoV-2 is a very contagious pathogen and the nasal viral loads in the asymptomatic patients are similar to that in the symptomatic patient, resulting in difficulties in controlling viral spread [5].

Small non-coding RNAs (sncRNAs) have diverse functions through various regulatory mechanisms. They virtually participate in all biological pathways, including cell proliferation, differentiation, apoptosis, autophagy, and tissue remodeling. sncRNAs are also essential to regulate host responses to viral infections [6-10]. Among sncRNAs, the most widely studied sncRNAs are microRNAs (miRNAs), which are 18∼24 nt in length, carry 5’ monophosphate and 3’hydroxyl (3’-OH) ends, and generally regulate genes via the argonaute (AGO) platform [11, 12]. Other than miRNAs, piwi-interacting RNAs (piRNAs), small nucleolar RNAs (snoRNAs), and tRNA-derived RNA fragments (tRFs) are also important members of sncRNAs [13]. Currently, there is very limited information on whether or how SARS-CoV-2 regulates the sncRNA expression, except the reports on SARS-CoV-2-impacted miRNAs [14-16].

Our recent seq data demonstrated that tRFs and piRNAs were the two most abundant sncRNAs in nasopharyngeal swabs (NPS) samples of the SARS-CoV-2-positive group. However, only tRFs, but not other types of sncRNAs, were significantly enhanced in SARS-CoV positive samples. Generally speaking, tRFs, ranging from 14∼40nt, are generated by specific cleavages within pre-tRNA or mature tRNAs [17]. Compared with other sncRNAs, tRFs are relatively new members. However, their importance in diseases, such as cancer, infectious diseases, neurodegenerative diseases, and metabolic diseases, was quickly acknowledged soon after the discovery [18-25]. They are generally classified into tRF-1 series, tRF-3 series, and tRF-5 series [26]. tRF-1 series are usually those from the 3′-trailer sequences of pre-tRNA, while tRF-3 and tRF-5 series are aligned to the 3′-and 5′-end of the mature tRNAs respectively. Among SARS-CoV-2-impacted tRFs, the most impacted tRFs belonged to tRF5s. In addition, the impacted tRF profile by SARS-CoV-2 seemed to be SARS-CoV-2 specific, which is consistent with what we and others found previously on the changes in tRF signatures being virus-dependent [21, 23], implicating tRFs as potential prognosis and diagnosis biomarkers. The impacted tRFs were also observed in SARS-CoV-2 infected human alveolar type II-like epithelial cells expressing human angiotensin-converting enzyme 2 (A549-ACE2) and human small airway epithelial cells (SAECs) in the air-liquid interface (ALI) culture.

In addition to host-derived ncRNAs, viral genomes can also encode ncRNAs. These viral ncRNAs vary in length and have diverse biological functions, including the regulation of viral replication, viral persistence, host immune evasion, host inflammatory response, and cell transformation [27]. For example, three18∼22 nt SARS-CoV-encoded small RNAs have been reported to contribute to SARS-CoV-induced lung injury [28]. Recently, multiple SARS-CoV-2-encoded miRNAs and their putative targets were first predicted by *in-silico* analysis by several independent research groups [29, 30]. The expression of some of the SARS-CoV-2-encoded miRNAs was later experimentally validated in NPS and lung tissues of COVID-19 patients and two SARS-CoV-2-encoded miRNAs could enhance the inflammation [31]. In this study, we also mapped our seq data to the viral genome and revealed several new small viral RNAs (svRNAs) fragments, with the length of 25 nt, 33 nt and 36 nt. Among CoV-2-derived svRNAs (sv-CoV2), sv-CoV2, derived from genomic site 346 to site 382 of nsp1 (sv-CoV2-346) had the highest expression. We confirmed its induction by RT-PCR and found that sv-CoV2-346 can only be detected when RNAs were pretreated with T4 PNK, demonstrating that svRNA lacking 3’-OH cannot be properly detected by routine small RNA seq or detected by unmodified RT-PCR.

In summary, this is the first report demonstrating the altered tRFs by SARS-CoV-2. T4 PNK pretreatment also enabled small RNA seq and RT-PCR to reveal additional new sv-CoV2 with a length longer than 22 nt. In the future, we will characterize the biogenesis and function mechanisms of these new sncRNAs associated with SARS-CoV-2 infection.

## Materials and methods

### Nasopharyngeal swabs (NPS) specimens

NPS were collected from patients who tested their SARS-CoV-2 in the out-patient clinics of the University of Texas Medical Branch (UTMB) in April 2020. NPS samples were transported to the Molecular Pathology laboratory, directed by Dr. Jianli Dong, in universal viral transport media, and subjected to SARS-CoV-2 test using Abbott m2000 SARS-CoV-2 RT-PCR assay. The limit of detection (LOD) of detection assays is 100 viral genome copies/ml.

Thirteen anonymous NPS samples were used in this study, including seven SARS-CoV-2 negative (51.7 ± 13.7 years old) and six SARS-CoV-2 positive (49.2 ± 10.5 years old) samples. The protocol was approved by institutional review boards (IRB) of UTMB at Galveston, under the IRB protocol # 02-089 and 03-385.

### RNA isolation

After the SARS-CoV-2 validation, 1 ml of NPS sample from each subject was subjected to RNA extraction using the mirVana PARIS kit (Invitrogen, MA, US) according to the manufacturer’s protocol. At the elution step, samples were incubated on the column for 5 min at 65°C, and the RNA was eluted with 45 µl nuclease-free water. RNAs of each sample were subjected to deep sequencing and qRT-PCR.

To extract RNAs from cells, TRIzol reagents (Thermo Fisher Scientific, MA, US) were used for total RNA preparation, as described [22], followed by qRT-PCR.

### T4 PNK-RNA-seq and data analyses

To study whether other sncRNAs than miRNAs are impacted by SARS-CoV-2, we used a modified next-generation sequencing (NGS), similar as described in [32] [33], to get sncRNA profiles for RNA samples, derived from NSP or cultured cells. Basically, we treated RNAs with T4 polynucleotide kinase (T4 PNK) before the library construction and small RNA-seq to make the 3’-end of RNAs homogenous with -OH, as the ligation of the 3’-end of sncRNAs with sequencing barcode requires the presence of 3’-OH modification and not all sncRNAs have such modification. We named T4 PNK-RNA-seq. The seq was done in the NGS Core of UTMB. In brief, the RNA samples were pretreated with 10 units of T4 PNK using 14 µl extracted RNAs in a final reaction volume of 50 µl and incubated at 37°C for 30 min, and then were heat-inactivated at 65°C for 20 min. The RNA was purified and concentrated within 6 µl nuclease-free water using Zymo RNA clean & concentrator kit (Irvine, CA, US). Ligation-based small RNA libraries were prepared with an RNA input of 6 µl using NEB Next Multiplex Small RNA Library Prep kit (Ipswich, MA, US). Libraries were sequenced using the Illumina NextSeq 550 Mid-Output sequencing run. About 7680 Mb of sequence data was generated.

Using cutadpat, adaptor sequences were removed and reads with a length of more than 15 bp were extracted. We further filtered out RNAs with counts of less than 10 and all rRNA sequences, using the remainders as cleaned input reads. In terms of the mapping databases, we prepared tRF5 and tRF3 databases using the same sequences derived from different tRNAs (sequences downloaded from tRNA genes using the Table Browser of the UCSC genome browser [34]). We also prepared tRF1 sequences using genome locations of tRNAs. Our in-house small RNA database includes 1) these tRFs, 2) miR/snoR sequences downloaded from the UCSC genome browser, and 3) piRNA sequences downloaded from piRBase (http://www.regulatoryrna.org/database/piRNA/). The cleaned input reads were mapped to our in-house small RNA database using bowtie2 (v2.4.1) allowing two mismatches (option m -1). After we mapped the cleaned input reads to the small RNA database, the unmapped sequences were then mapped to the hg38 genome using the bowtie2 pre-built index (GRCh38_noalt_as) to detect all human sequences. The unmapped sequences to the human genome were then mapped to the SARS-CoV-2 reference genome (NC_045512) using the same parameters.

Raw read counts were normalized with the DEseq2 median of ratios method. Differentially expressed genes were determined by p-value < 0.05, fold change >2, and read basemean > 10 in either CN or SARS-CoV-2 group. Unsupervised hierarchical clustering was performed using the Pearson correlation coefficient.

### Cell culture and viruses

African green monkey kidney epithelial cells (Vero E6) were obtained from ATCC (Manassas, VA, US), and maintained in a high-glucose DMEM (Gibco, MA, US) supplemented with 10% fetal bovine serum (FBS), 10 units/ml penicillin, and 10 µg/ml streptomycin. The human alveolar type II-like epithelial cells expressing human angiotensin-converting enzyme 2 (A549-ACE2) cells were a kind gift from Dr. Shinji Makino and were cultured in DMEM (Gibco, MA, US) containing 10% FBS, 10 units/ml penicillin, and 10 µg/ml streptomycin.

Small airway epithelial cells (SAECs), isolated from the normal distal portion of the lung in the 1 mm bronchiole area, were purchased from Lonza (Basel, Switzerland) to generate cells in the air-liquid interface (ALI) culture. The cells were cultured and differentiated using Complete PneumaCult™-Ex plus medium and PneumaCult™-ALI-S Maintenance medium (Stemcell Technologies, Vancouver, Canada), respectively, according to the manufacturer’s instructions. Briefly, the cells at passage two (P2) were expanded in the T-25 flask using the Complete PneumaCultTM-Ex plus medium, with a medium change every other day. For ALI cultures, the cells (P3) were seeded into Corning Costar 12 mm transwell inserts (Corning, NY, US) at a concentration of 11× 10^4^ cells/insert in 0.5 ml medium/insert, and another 1 ml/well medium was added to the basal chamber. Cells were submerged cultured in Complete PneumaCultTM-Ex plus medium, with a medium change every other day. After reaching ∼100% confluency, ALI was initiated by removing the apical medium and replacement of the PneumaCultTM-Ex plus medium in the basal compartment with PneumaCult™-ALI-S Maintenance medium. The basal compartment medium was changed every other day. It took about 21 days to complete cell differentiation.

SARS-CoV-2 (USA-WA1/2020 strain) was obtained from the World Reference Center for Emerging Viruses and Arboviruses (WRCEVA) at the UTMB. Viral stocks were prepared by propagation in Vero E6 cells. Viral titers were determined by plaque assay as described [35]. All experiments using live SARS-CoV-2 were performed in a biosafety level 3 (BSL-3) laboratory with redundant fans in the biosafety cabinets. All personnel wore powered air-purifying respirators (Breathe Easy, 3M) with Tyvek suits, aprons, booties, and double gloves.

### Viral infection

To infect A549-ACE2 cells in monolayer culture, the cells were seeded into the 24-well plate 24 h prior to the infection to allow the cells to reach 80∼90% confluence in the following day. For infection, the cells were incubated with viruses in DMEM media with 10% FBS at a multiplicity of infection of 0.1 (MOI = 0.1). After 1 h incubation, cells were washed with PBS for three times to remove the remaining viruses and cultured in fresh media containing 10% FBS. The cells were collected on day 4 post-infection. Three independent biological replicates were performed.

Regarding the infection of SAECs in ALI culture, the infection was performed when hallmarks of excellent differentiation were evident, such as extensive apical coverage with cilia. Prior to infection, the apical side of the cells was washed five times with PBS, and the basal surface was washed once with PBS. Viruses were diluted to the specified MOI in 200 µl MEM medium and inoculated onto the apical surface of the ALI cultures. After a 2-hour incubation at 37 °C with 5% CO2, unbound viruses were removed by washing with PBS three times. The cells were collected on day 1 or 3 post-infection. The SARS-CoV-2 S gene was detected using qRT-PCR with primers as follows: S forward primer, 5’ CCTACTAAATTAAATGATCTCTGCTTTACT; reverse primer, 5’ CAAGCTATAACGCAGCCTGTA.

### qRT-PCR and RT-PCR

To evaluate sncRNAs expression, qRT-PCR was performed as described previously [22, 24]. Briefly, the total RNA was treated with T4 PNK, and then ligated to a 3’-RNA linker using T4 RNA ligase from Thermo Fisher Scientific (Waltham, MA, US). The product was used as a template for reverse transcription (RT) with a linker-specific reverse primer using TaqMan Reverse Transcription Reagents from Thermo Fisher Scientific. The cDNA was subjected to SYBR Green qPCR using iTaq™ Universal SYBR Green Supermix kit from Bio-Rad (Hercules, CA, US) and primers specific to the 5’-end of tRFs and RNA linker. U6 was used for normalization. The addition of a 3’-RNA linker enables the detection of tRF5s without the interference of its corresponding parent tRNAs. The sequences of the primers and the 3’-RNA linker are listed in **Table I**.

**Table I.**
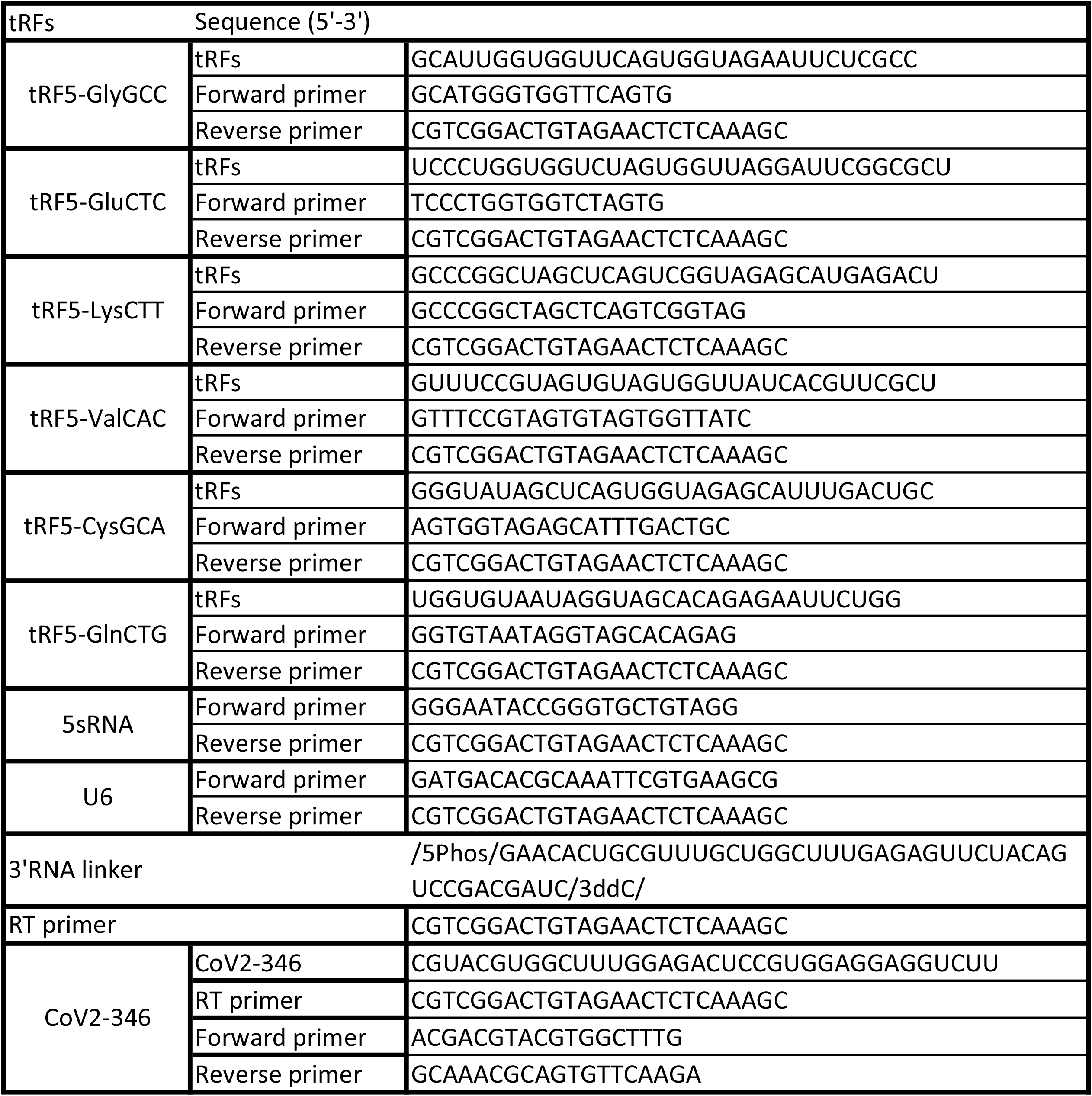
Sequence information of tRF5s, RNA linker, RT primer, and PCR/qPCR primers (experimental validation)

To validate the seq data on the presence of CoV2-encoded small RNAs (CoV2-346), RT-PCR was performed, using RT and PCR primers listed in **Table I**. Briefly, RNAs, pretreated with or without T4 PNK, were also ligated to the 3’RNA linker. The RT was done using primers complementary to the RNA linker, followed by PCR using forward primers annealing to 5’-end of CoV2-346 and reverse primers annealing to the last 4 nt of CoV2-346 and the first 15 nt of RNA-linker.

### Northern blot

Northern hybridization for tRF5-GluCTC was performed as described [21]. Briefly, 3 µg RNA was loaded on 15% denaturing polyacrylamide gel with 7 M ureas and then transferred to a positively charged nylon membrane (Amersham Biosciences, NJ, US). The membrane was hybridized with a ^32^P-labeled DNA probe in ULTRAhyb-Oligo solution (Life Technologies, Grand Island, NY, US) and washed three times according to the manufacturer’s instruction, followed by image development.

### Statistical analysis

The experimental results were analyzed using Graphpad Prism 5 software. To compare the sncRNAs expression of NPS between SARS-CoV-2 negative and positive groups, an unpaired two-tailed Mann-Whitney U test was used. To compare the sncRNAs expression in SARS-CoV-2 infected cells and mock-infected cells, an unpaired two-tailed t-test was employed. A *p*-value < 0.05 was considered to indicate a statistically significant difference. Single and two asterisks represent a p-value of < 0.05 and <0.01, respectively. Means ± standard errors (SE) are shown.

## Results

### T4 PNK-RNA-seq revealed SARS-CoV-2 impacted sncRNAs in NPS samples

To identify SARS-CoV-2-impacted sncRNAs, we initialized T4 PNK-RNA-seq for the NPS samples from four SARS-CoV-2 positive patients with their ages at 54.3 ± 4.0 years old and four SARS-CoV-2 negative subjects, with matched age at 50.5 ± 10.2 years old. The seq data were analyzed similarly as described in [21, 36]. In brief, the sequences with length > 15 bp and reads > 10 were mapped to the in-house small RNA database containing tRFs, miR/snoRs, and piRs to address redundant tRNA sequences across the genome after removing rRNAs. Unmapped sequences were then mapped to the hg38 human genome to identify other human-derived sequences and their composition. We found that piRNAs and tRFs were the two most abundant sncRNAs in SARS-CoV-2 positive samples. The top-10 ranked piRNAs and tRFs in the SARS-CoV-2 positive group are listed in **Supplementary Tables I** and **II**, respectively. As shown in **Supplementary Table II**, all tRFs were derived from the 5’-ends of tRNAs, therefore tRF5s. Compared with the tRFs and piRNAs, the overall reads of miRNAs were much less (**Figure 1A**). We also compared the sncRNA profiles between SARS-CoV-2 positive and negative samples. As shown in **Figure 1A**, while the tRFs consisted of about 14% of all sncRNA counts in the control group, tRFs counts became 42% in the COVID-19 group, demonstrating a significant increase by COVID-19. In contrast, the overall expression of miRNAs and piRNAs was not impacted by COVID-19 (**Figure 1A**).

**Figure 1.**
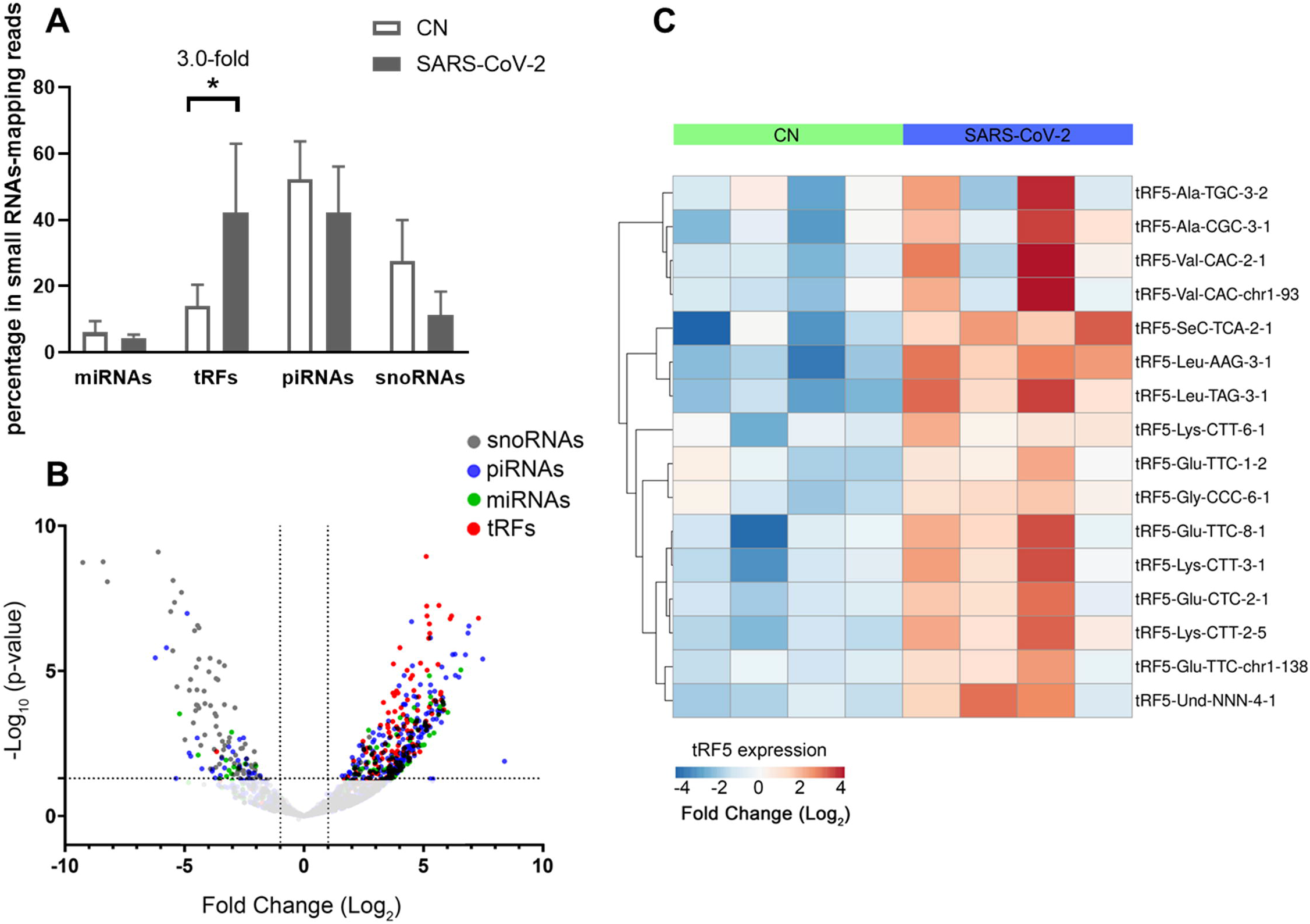
Impacted sncRNAs by SARS-CoV-2 infection in patients. **(A)** The relative sequencing frequency of miRNAs, tRFs, piRNAs, and snoRNAs was calculated by normalizing their raw reads with the DEseq2 median ratio method. **(B)** The volcano plot showed that sncRNAs were differentially expressed between and control group (CN) and SARS-CoV-2 patient group (SARS-COV-2). **(C)** Heatmap for unsupervised clustering of the differently expressed tRFs with > 20 basemean according to Pearson correlation. Data are shown as means ± standard error (SE). The single asterisk represents *P* values of <0.05.

Differential expression analysis for individual sncRNAs was also performed for SARS-CoV-2 negative and positive groups. As shown in **Figure 1B**, there were more up-regulated tRFs than down-regulated tRFs, while SARS-CoV-2 down-regulated snoRNAs were more than up-regulated ones. We also listed sncRNAs, which were significantly altered by SARS-CoV-2 **in Tables II-IV**. The cutoff was set as a fold change >2, with the significance of *p* < 0.05 in changes by SARS-CoV-2, and the basemean > 10 in the negative or positive group. The differentially expressed tRFs, miRNAs, and snoRNAs were listed in **Table II, Table III, and Table IV**, respectively.

**Table II.**
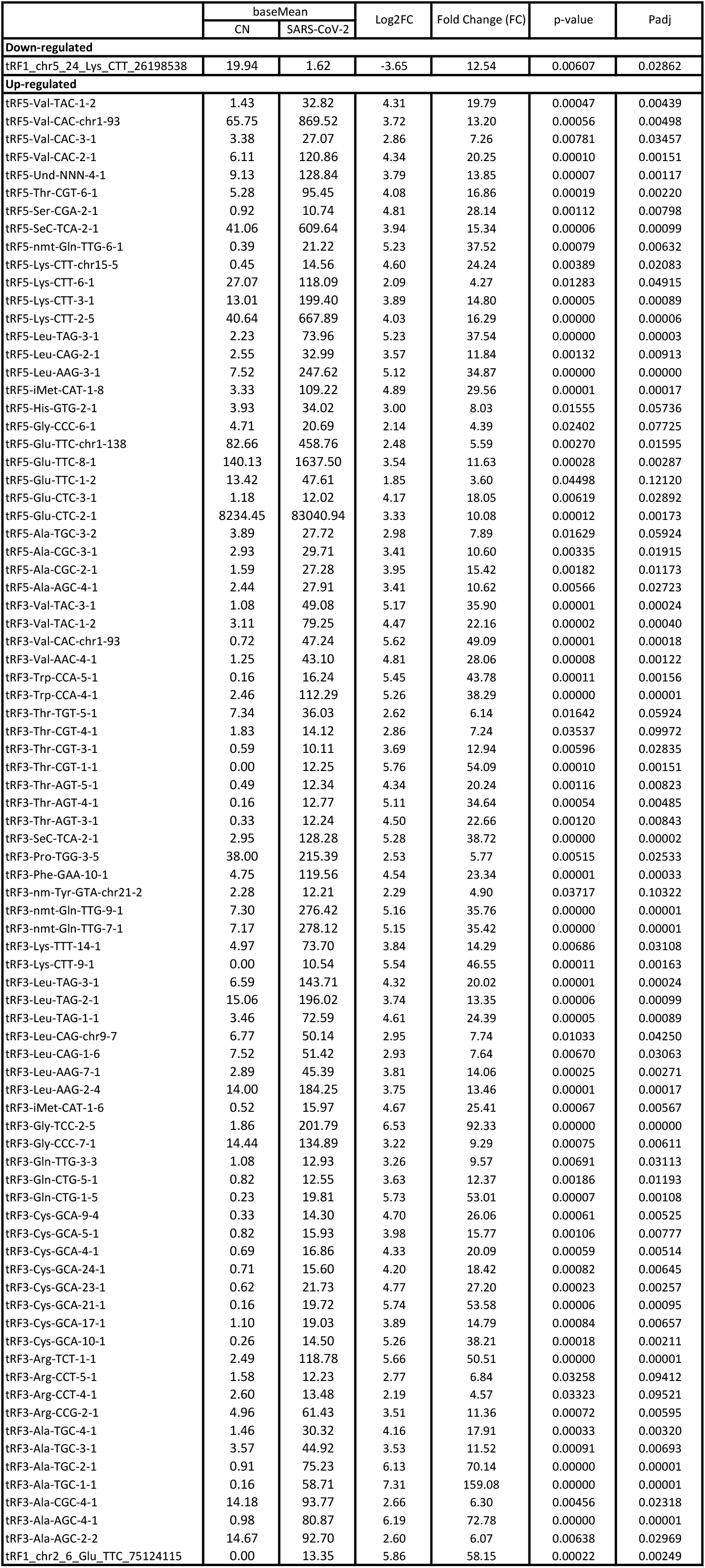
SARS-CoV-2-impacted tRFs

**Table III.**
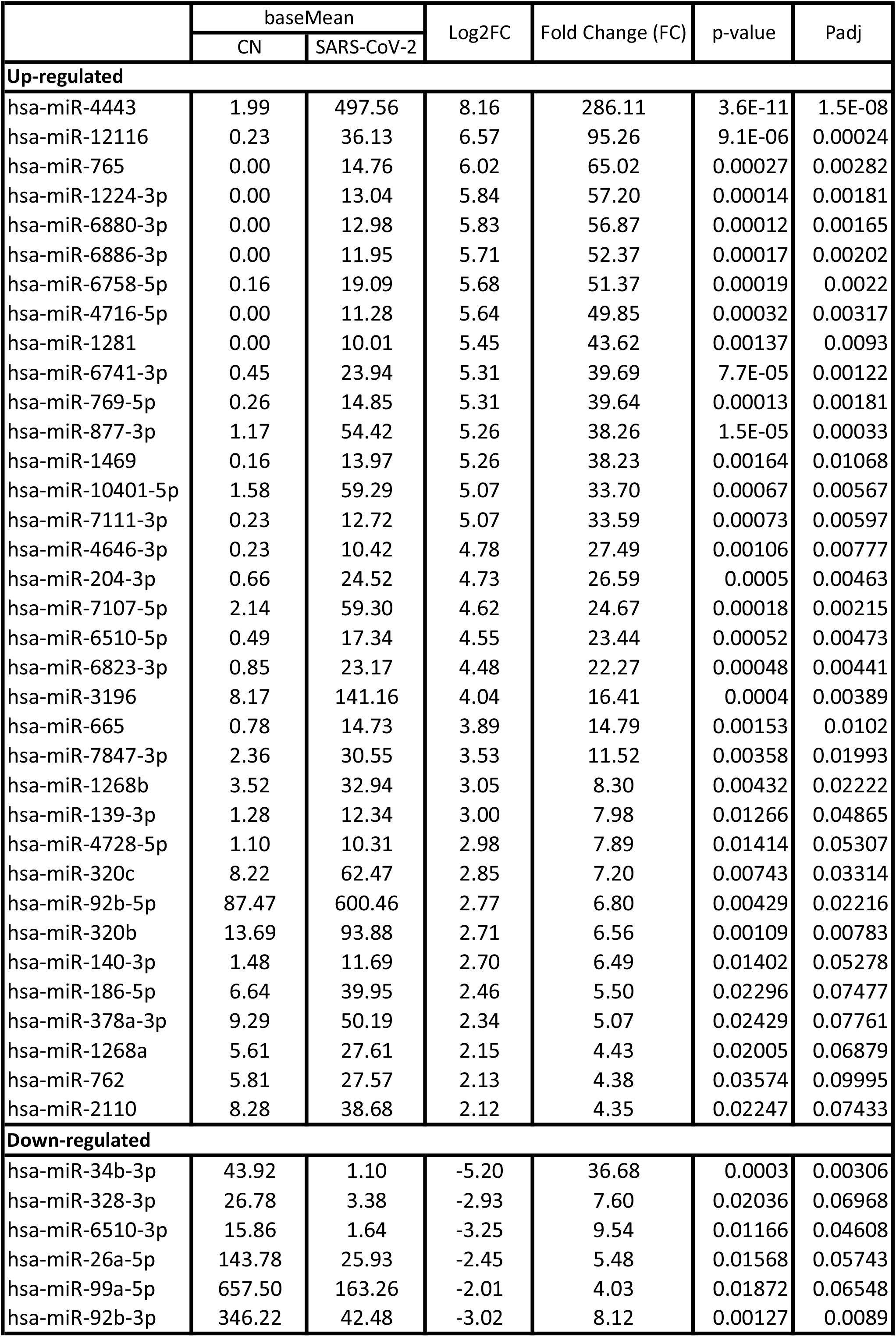
SARS-CoV-2-affected miRNAs.

**Table IV.** SARS-CoV-2-changed snoRNAs and their related information.

|  | baseMean |  | Log2FC | Fold Change (FC) | p-value | Padj |
| --- | --- | --- | --- | --- | --- | --- |
|  | CN | SARS-CoV-2 |  |  |  |  |
| Down-regulated |  |  |  |  |  |  |
| hg38_wgRna_ACA49 | 227.26 | 0.27 | -9.25 | 609.86 | 1.9E-09 | 3.4E-07 |
| hg38_wgRna_ACA40 | 180.98 | 0.53 | -8.40 | 338.05 | 1.7E-09 | 3.4E-07 |
| hg38_wgRna_ACA28 | 219.23 | 0.72 | -8.22 | 299.21 | 8.6E-09 | 1.1E-06 |
| hg38_wgRna_ACA8 | 409.75 | 8.53 | -5.57 | 47.58 | 9.1E-08 | 7.7E-06 |
| hg38_wgRna_U92 | 79.97 | 1.72 | -5.49 | 44.82 | 2.1E-06 | 8E-05 |
| hg38_wgRna_E3 | 162.73 | 3.73 | -5.41 | 42.66 | 4.4E-08 | 4.7E-06 |
| hg38_wgRna_ACA25 | 38.67 | 0.92 | -5.31 | 39.75 | 3.6E-05 | 0.00069 |
| hg38_wgRna_ACA16 | 158.54 | 4.34 | -5.13 | 34.92 | 2E-08 | 2.3E-06 |
| hg38_wgRna_ACA3 | 206.55 | 7.70 | -4.74 | 26.71 | 2E-05 | 0.00042 |
| hg38_wgRna_U73a | 25.28 | 1.01 | -4.62 | 24.63 | 0.00062 | 0.00531 |
| hg38_wgRna_U35A | 99.03 | 4.05 | -4.58 | 23.87 | 4.2E-07 | 2.1E-05 |
| hg38_wgRna_ACA38 | 35.59 | 1.45 | -4.53 | 23.09 | 0.00024 | 0.00258 |
| hg38_wgRna_mgU6-77 | 93.30 | 4.01 | -4.50 | 22.65 | 7.6E-06 | 0.0002 |
| hg38_wgRna_HBII-82B | 36.20 | 1.76 | -4.50 | 22.57 | 0.00015 | 0.00187 |
| hg38_wgRna_E2 | 88.91 | 4.05 | -4.48 | 22.26 | 4.5E-05 | 0.00087 |
| hg38_wgRna_ACA63 | 122.12 | 5.51 | -4.45 | 21.84 | 0.00016 | 0.00192 |
| hg38_wgRna_HBII-180A | 157.02 | 7.50 | -4.39 | 21.04 | 4E-06 | 0.00013 |
| hg38_wgRna_U71a | 38.72 | 1.83 | -4.39 | 20.95 | 0.00019 | 0.00221 |
| hg38_wgRna_ACA41 | 21.63 | 1.00 | -4.31 | 19.86 | 0.00372 | 0.02024 |
| hg38_wgRna_U90 | 86.50 | 4.66 | -4.18 | 18.07 | 5E-05 | 0.00089 |
| hg38_wgRna_ACA10 | 38.56 | 2.01 | -4.15 | 17.73 | 0.00162 | 0.0106 |
| hg38_wgRna_U97 | 89.21 | 4.93 | -4.14 | 17.68 | 0.00013 | 0.00181 |
| hg38_wgRna_ACA44 | 114.67 | 7.12 | -3.97 | 15.72 | 1.1E-05 | 0.00027 |
| hg38_wgRna_ACA9 | 13.58 | 1.00 | -3.75 | 13.48 | 0.00509 | 0.02516 |
| hg38_wgRna_ACA51 | 85.91 | 6.42 | -3.72 | 13.16 | 0.0003 | 0.00306 |
| hg38_wgRna_ACA3-2 | 124.34 | 9.63 | -3.68 | 12.78 | 0.00082 | 0.00645 |
| hg38_wgRna_U15A | 87.31 | 6.73 | -3.67 | 12.73 | 0.00036 | 0.00352 |
| hg38_wgRna_U15B | 37.76 | 3.37 | -3.53 | 11.57 | 0.00156 | 0.01027 |
| hg38_wgRna_ACA1 | 28.02 | 2.69 | -3.43 | 10.81 | 0.00929 | 0.0395 |
| hg38_wgRna_ACA53 | 62.15 | 5.85 | -3.41 | 10.60 | 0.0027 | 0.01595 |
| hg38_wgRna_SNORD123 | 21.14 | 2.10 | -3.36 | 10.30 | 0.00525 | 0.02556 |
| hg38_wgRna_U17b | 86.15 | 8.67 | -3.32 | 10.02 | 0.00015 | 0.00184 |
| hg38_wgRna_ACA6 | 31.12 | 3.13 | -3.31 | 9.89 | 0.00373 | 0.02024 |
| hg38_wgRna_U34 | 33.68 | 3.92 | -3.15 | 8.87 | 0.00286 | 0.01669 |
| hg38_wgRna_U18B | 16.65 | 1.90 | -3.07 | 8.38 | 0.01113 | 0.04479 |
| hg38_wgRna_SNORA38B | 13.91 | 1.82 | -2.91 | 7.49 | 0.01566 | 0.05743 |
| hg38_wgRna_U32A | 70.17 | 9.50 | -2.86 | 7.27 | 0.00813 | 0.03551 |
| hg38_wgRna_U52 | 41.92 | 6.17 | -2.79 | 6.92 | 0.03576 | 0.09995 |
| hg38_wgRna_U51 | 26.73 | 4.02 | -2.75 | 6.73 | 0.02161 | 0.07241 |
| hg38_wgRna_U17a | 28.75 | 4.49 | -2.64 | 6.21 | 0.00479 | 0.02396 |
| hg38_wgRna_HBII-85-18 | 38.90 | 6.61 | -2.53 | 5.76 | 0.02759 | 0.08448 |
| hg38_wgRna_U19-2 | 18.33 | 4.68 | -2.00 | 4.00 | 0.04758 | 0.12608 |

As shown in **Table II**, tRFs were significantly impacted by SARS-CoV-2 in NPS. Among those, 2 tRFs belong to the tRF1 series, 28 tRFs were tRF5s, and 53 tRFs were tRF3s. However, top-ranked SARS-CoV-2-impacted tRFs all belong to the tRF5. In **Figure 1C**, the read basemean > 20 in control (CN) or SARS-CoV-2 positive (SARS-CoV-2) group were selected to plot the heatmap and their sequences were listed in **Table V**.

**Table V.**
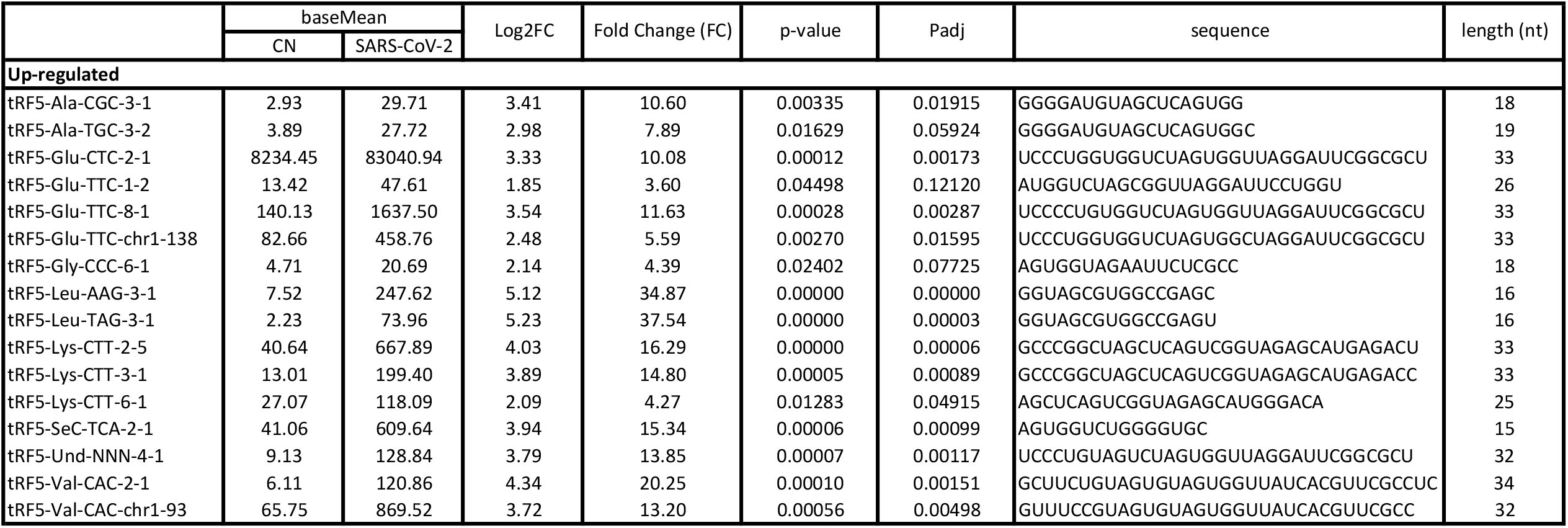
SARS-CoV-2-impacted tRF5, with basemean > 20 in contorl (CN) and/or SARS-CoV-2 postive (SARS-COV-2) group.

### Experimental validation of SARS-CoV-2-impacted tRFs

To validate the seq data, we used modified qRT-PCR to detect expression of tRF5-GluCTC, tRF5-LysCTT, and tRF5-ValCAC, three top-ranked tRF5s in SARS-CoV-2-positive patients and also SARS-CoV-2-impacted tRFs according to the seq data, as we previously described [22, 24]. Compared with Northern blot validation, the modified qRT-PCR with T4 PNK pretreatment and 3’-end RNA linker ligation provides the possibility to validate as many tRFs as possible for NPS samples, which usually have a limited amount of samples, therefore, the limited yield of RNAs. Our results demonstrated that tRF5-GluCTC, tRF5-LysCTT, and tRF5-ValCA were significantly increased in the SARS-CoV-2 group (**Figures. 2A-C**). Unlike SARS-CoV-2, which could induce tRF5-ValCAC, respiratory syncytial virus (RSV), a negative-sense RNA virus, does not induce tRF5-ValCAC infected cells [21], suggesting the change in tRF profile in response to viral infections is virus-specific.

**Figure 2.**
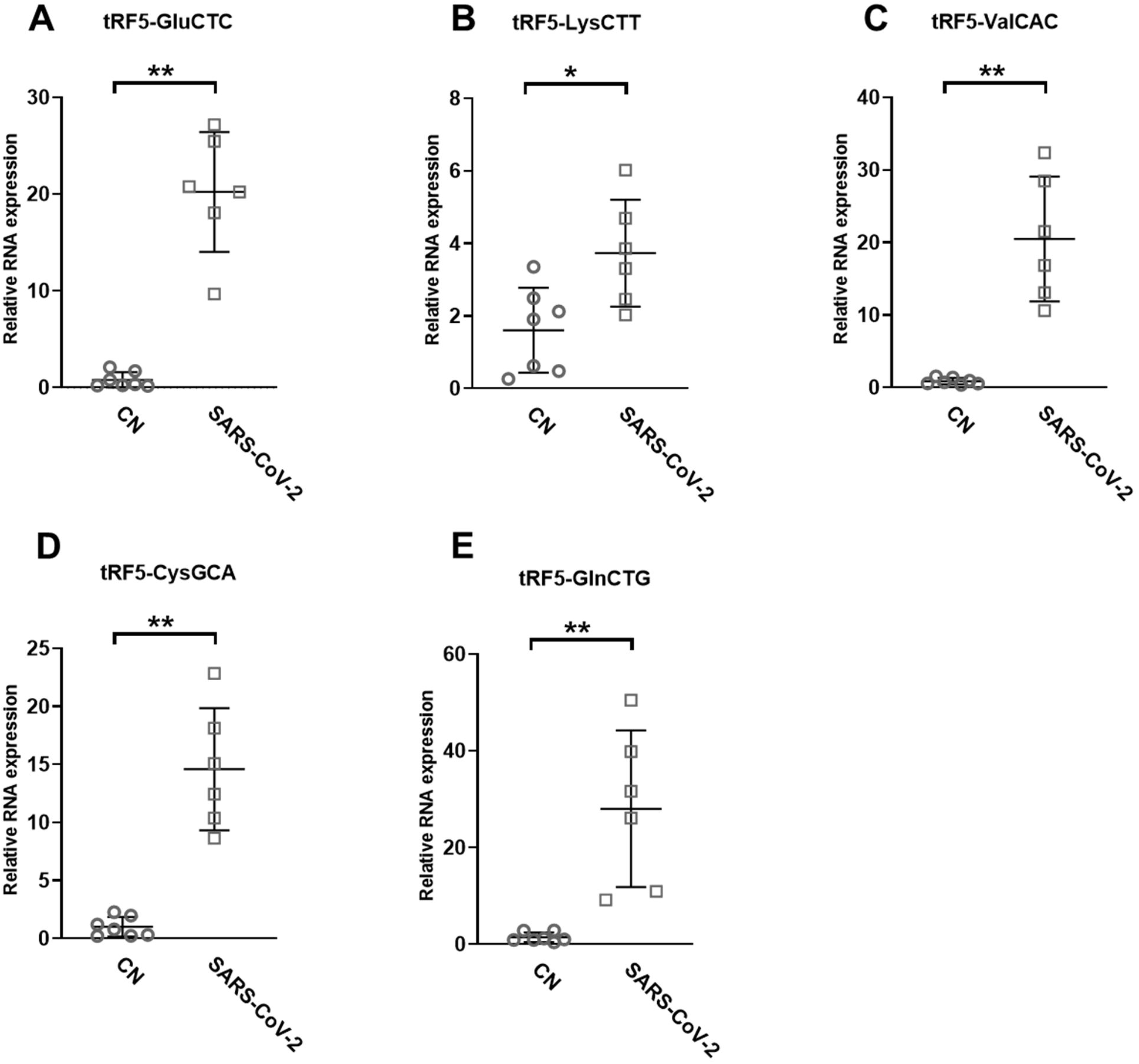
Changes in the expression of tRF5s in NPS samples by SRAS-CoV-2. (A-F) qRT-PCR was performed to detect tRF5-GluCTC **(A)**, tRF5-LysCTT **(B)**, tRF5-ValCAC **(C)**, tRF5-CysGCA **(D)**, and tRF5-GlnCTG **(E)** in the NPS from SARS-COV-2 and control (CN) patients. U6 was used as an internal control. Unpaired two-tailed Mann-Whitney U tests were performed for statistical comparisons. Single and double asterisks represent *P* values of <0.05 and <0.01, respectively. Data are shown as means ± SE.

Other than the three tRF5s mentioned above, tRF5-CysGCA and tRF5-GlnCTG were also chosen for the validation, as these two tRFs are highly inducible by RSV with the function tRF5-GlnCTG being proviral [22, 37]. Although the function of tRF5-CysGCA is not clear in viral infection, it is functionally important in stress responses and neuroprotection [38]. We validated that these two tRF5s were also significantly enhanced in the SARS-CoV-2 group, compared with the CN group (**Figures 2D and E**).

### Impacted tRFs in SARS-CoV-2-infected cells

A549-ACE2 is a wildly used cell model to study coronaviruses, such as SARS-CoV and SARS-CoV-2 [39-42]. Herein, we studied whether the induction of tRFs can be recapitulated in SARS-CoV-2-infected A549-ACE2. As shown in **Figures 3A-E**, A549-ACE2 cells, at day 4 post-infection (p.i.) of SARS-CoV-2 at an MOI of 0.1, had dramatically induction of tRF5-GluCTC, tRF5-LysCTT, tRF5-ValCAC, tRF5-CysGCA, and tRF5-GlnCTG. The northern blot also confirmed the induction of tRF5-GluCTC (**Figure 3F**), which was the most abundant tRF5 among the tested four tRF5s, confirming the liability of our newly developed qRT-PCR.

**Figure 3.**
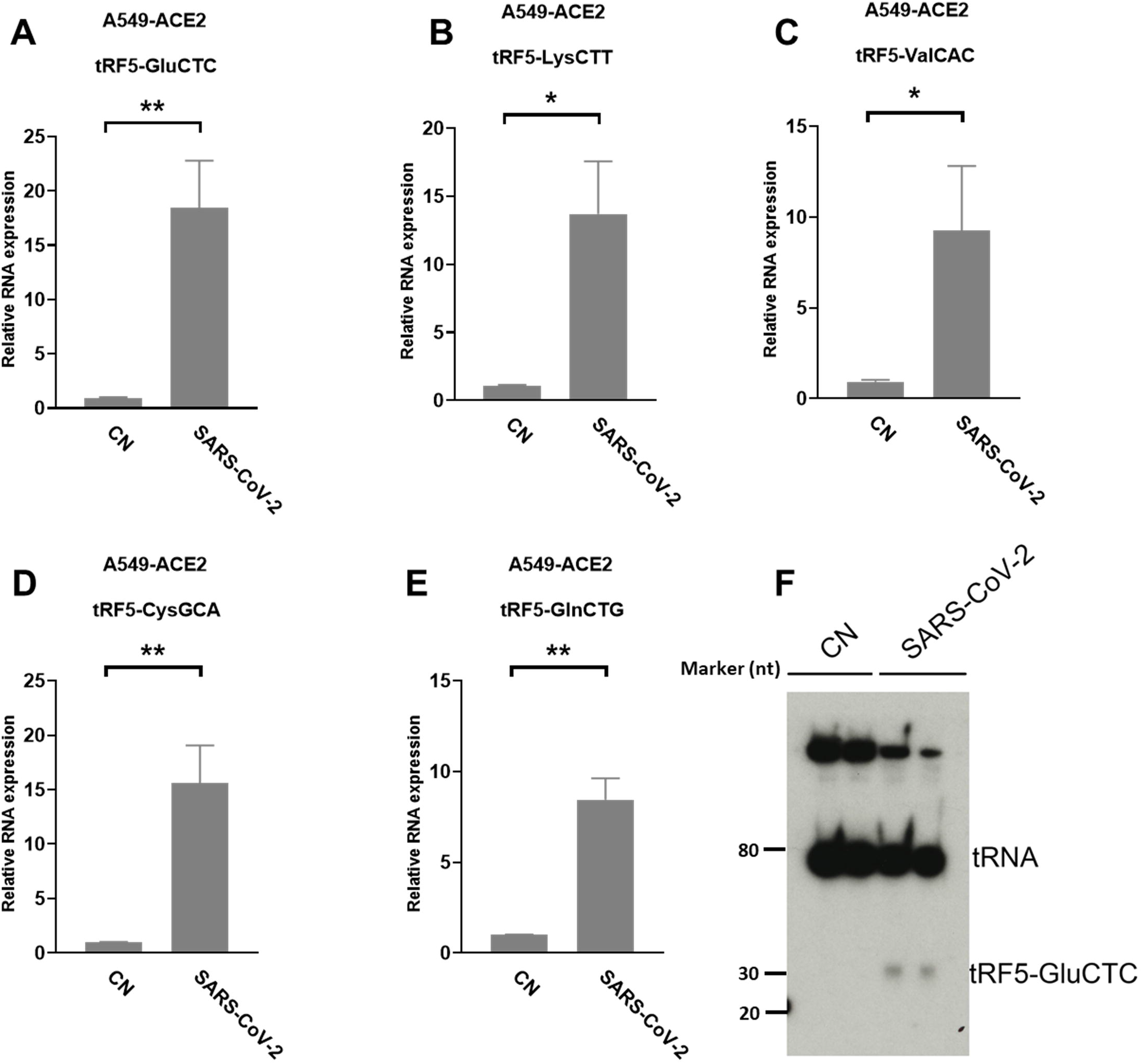
SARS-CoV-2-impacted tRF5s in A549-ACE2 cells. **(A-E)** A549-ACE2 cells were infected with SARS-CoV-2 at an MOI of 0.1 for 4 days, qRT-PCR was performed to detect tRF5-GluCTC **(A)**, tRF5-LysCTT **(B)**, tRF5-ValCAC **(C)**, tRF5-CysGCA **(D)**, and tRF5-GlnCTG **(E)**. A northern blot was performed to detect tRF5-GluCTC **(F)**. Unpaired two-tailed t-tests were performed for statistical comparisons. Single and double asterisks represent *P* values of <0.05 and <0.01, respectively. Data, shown as means ± SE, are representative of three independent experiments.

We also used primary SAECs in ALI culture, a commonly acknowledged physiology airway infection model[43, 44], to confirm SARS-CoV-2-affected tRFs. SAECs, after a few weeks of ALI culture, have been shown to establish a pseudostratified cell layer that resembles the small airway epithelium as found *in vivo* [45]. Moreover, SAECs in ALI cultures have been found to express the receptor for SARS-CoV-2, therefore, also a cell model to study SARS-CoV-2 [1, 46, 47]. Therefore, we also studied tRF5s expression in SAECs in ALI culture. As shown in **Figures 4A and B**, the cilia were oriented towards the upper transwell compartment, after the cells were in ALI culture for 21 days. The differentiated cultures were infected with SARS-CoV-2 at an MOI of 0.1 for 1 or 3 days, followed by viral S gene quantification using qRT-PCR (**Figure 4C**). Our qRT-PCR revealed that the expression change in tRF5-GlnCTG and tRF5-ValCAC, two tRFs with relatively low abundance in SARS-CoV-2 positive NPS samples and infected A549-ACE2 cells, can also be detected in SAECs in ALI culture (**Figures 4 D and E**). Overall, in this study, we established two cell models, A549-AEC2 cells in monolayer culture and SAECs in ALI culture, to characterize SARS-CoV-2-induced tRFs in the future.

**Figure 4.**
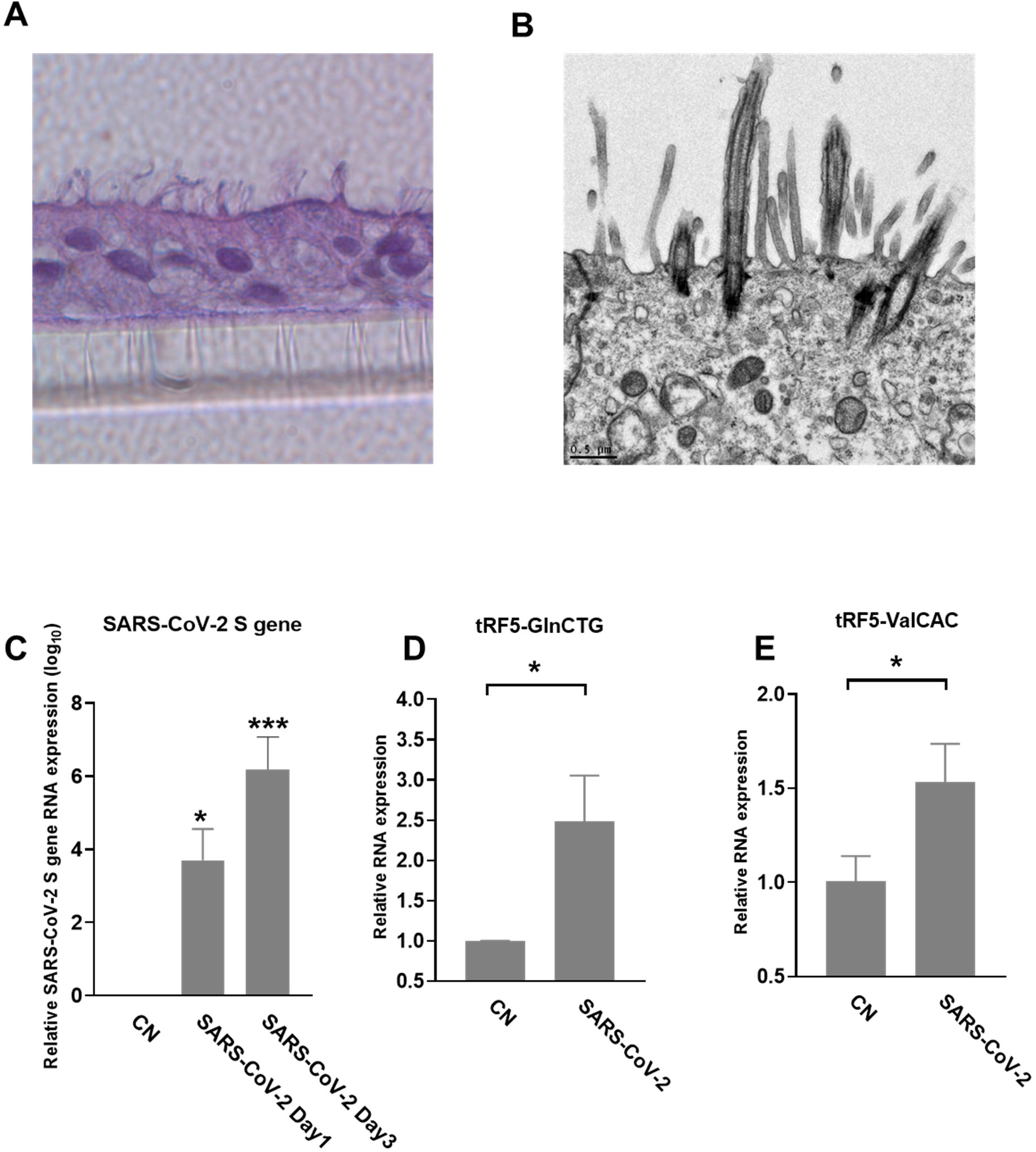
SARS-CoV-2-affected tRF5s in ALI-cultured SAECs. **(A)** Histological examination SAECs in ALI culture. After SAECs were in ALI culture for 21 days, the insert was fixed with 4% polyoxymethylene, followed by tissue processing, sectioning, and H&E staining. **(B)** Transmission electron microscopy (TEM) of SAECs in ALI culture. **(C)** ALI-cultured SEACs were apically infected with SARS-CoV-2 at an MOI of 0.1 for 1 or 3 days, qRT-PCR was performed to detect SARS-CoV-2 S gene expression. GAPDH was used as an internal control. **(D-E)** On day 3 post-infection, tRF5-GlnCTG (D) and tRF5-ValCAC (E) expressions were measured using qRT-PCR. Unpaired two-tailed t-tests were performed for statistical comparisons. Single asterisks represent *P* values of <0.05. Data are shown as means ± SE and are representative of three independent experiments.

### Identification of SARS-CoV-2-encoded svRNAs

Viral-derived sncRNAs are also an important family of sncRNAs [27]. To investigate whether SARS-CoV-2-encoded svRNAs are produced in the context of SARS-CoV-2 infection, the seq data were also aligned to the complete genome sequence of SARS-CoV-2 isolate Wuhan-Hu-1 (NC_045512.2). Several SARS-CoV-2-derived svRNAs were identified in SAARS-CoV-2 positive samples. The eight most abundant SARS-CoV-2-encoded svRNAs are listed in **Table. VI**.

**Table VI.**
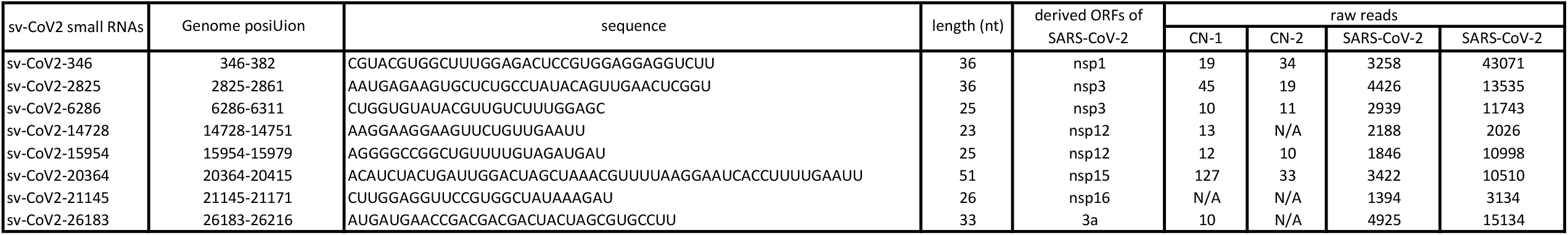
SARS-CoV-2-encoded svRNAs and their related information.

We further analyzed the sequences of svRNAs. Since only RNA <200 bp were selected for cDNA library, our results should not give scRNAs larger than 200 bp. Our results indicated the length of mapped svRNAs ranged from 17 to 75 nt, Among the svRNAs, svRNAs with the length of 25 nt, 33 nt, and 36 nt were enriched (**Figure. 5A**, two representatives are shown). Overall the enriched svRNAs are longer than the canonical miRNAs. In **Figure 5B**, the locations of the top 8 svRNAs along the SARS-CoV-2 genome are shown.

**Figure 5.**
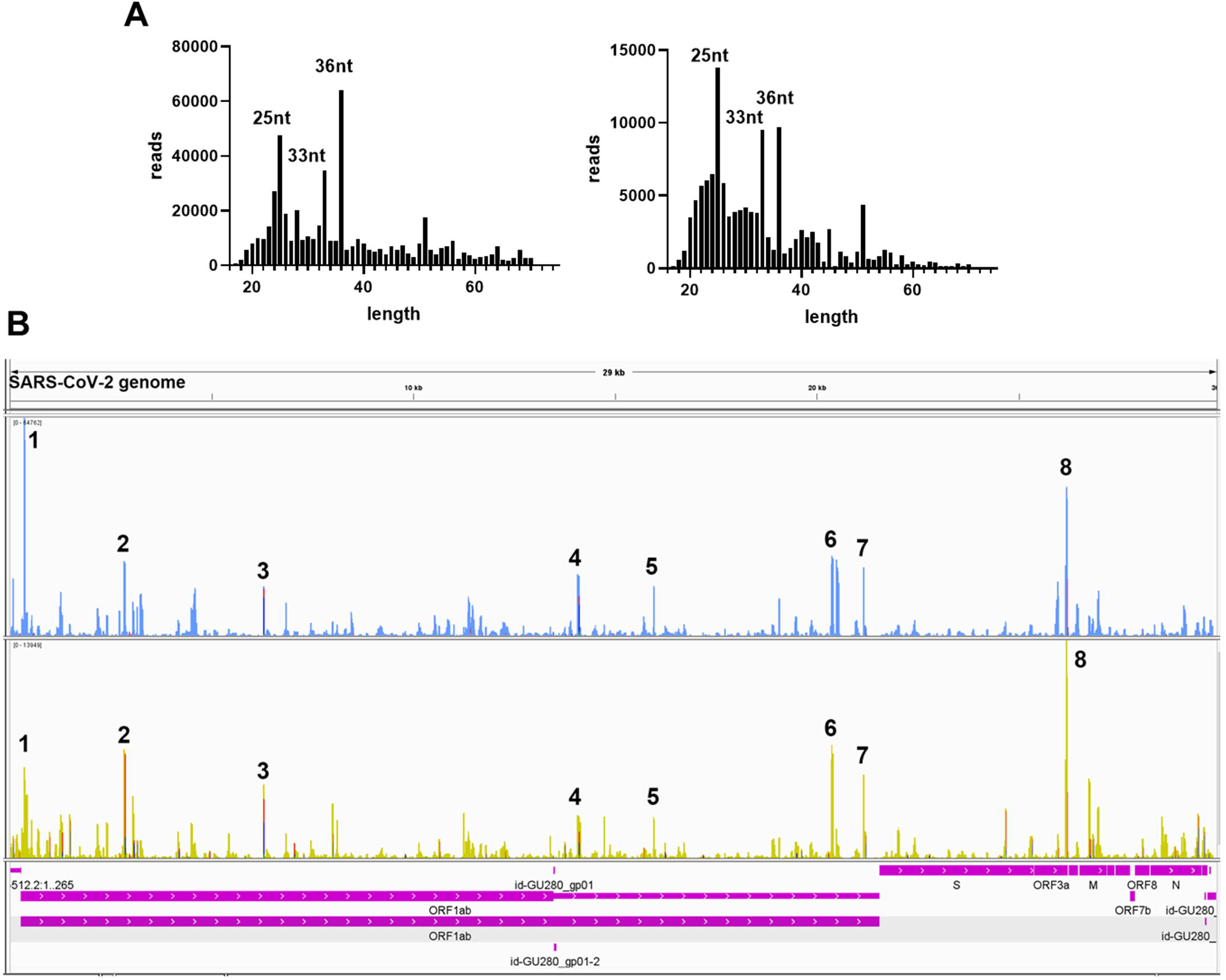
Viral small RNA sequence information. **(A)** viral small RNA sequence length distribution of two representative patient samples. **(B)** Two representative visual inspections of the small RNA sequences aligning with the SARS-CoV-2 genome. The names of viral genes and the genome positions (nt) are indicated.

### Experimental validation of SARS-CoV-2-encoded svRNAs

Among CoV-2-derived svRNAs (sv-CoV2), a 36 nt sv-CoV2, derived from genomic site 346 to site 382 of nsp1 (sv-CoV2-346) had the highest expression. To further validate the presence of sv-CoV2-346, NPS RNAs from two control samples and two COVID-19 samples were treated with T4 PNK and linked to a 3’ RNA adaptor, and then the RT-PCR was performed. The RT-PCR was also performed without the T4 PNK treatment and RNA linker addition so that the importance of such treatments can be demonstrated. The overall workflow is illustrated in **Figures 6A and B**. The specific 55nt RT-PCR products of sv-CoV2-346 were observed in SARS-CoV-2 samples, but not in the control samples, when samples were pretreated with T4 PNK and ligated with an RNA linker (**Figure. 6C**). The length reflected the 36 nt sv-CoV2-346 along with the 3’ RNA linker. In addition, we found that the RT-PCR product of one SARS-CoV-2 sample was more than another one, which was consistent with their read frequence in Seq-data. The presence of sv-CoV2-346 was confirmed in SARS-CoV-2-infected A549-ACE2 cells using RT-PCR (**Figure. 6D**).

**Figure 6.**
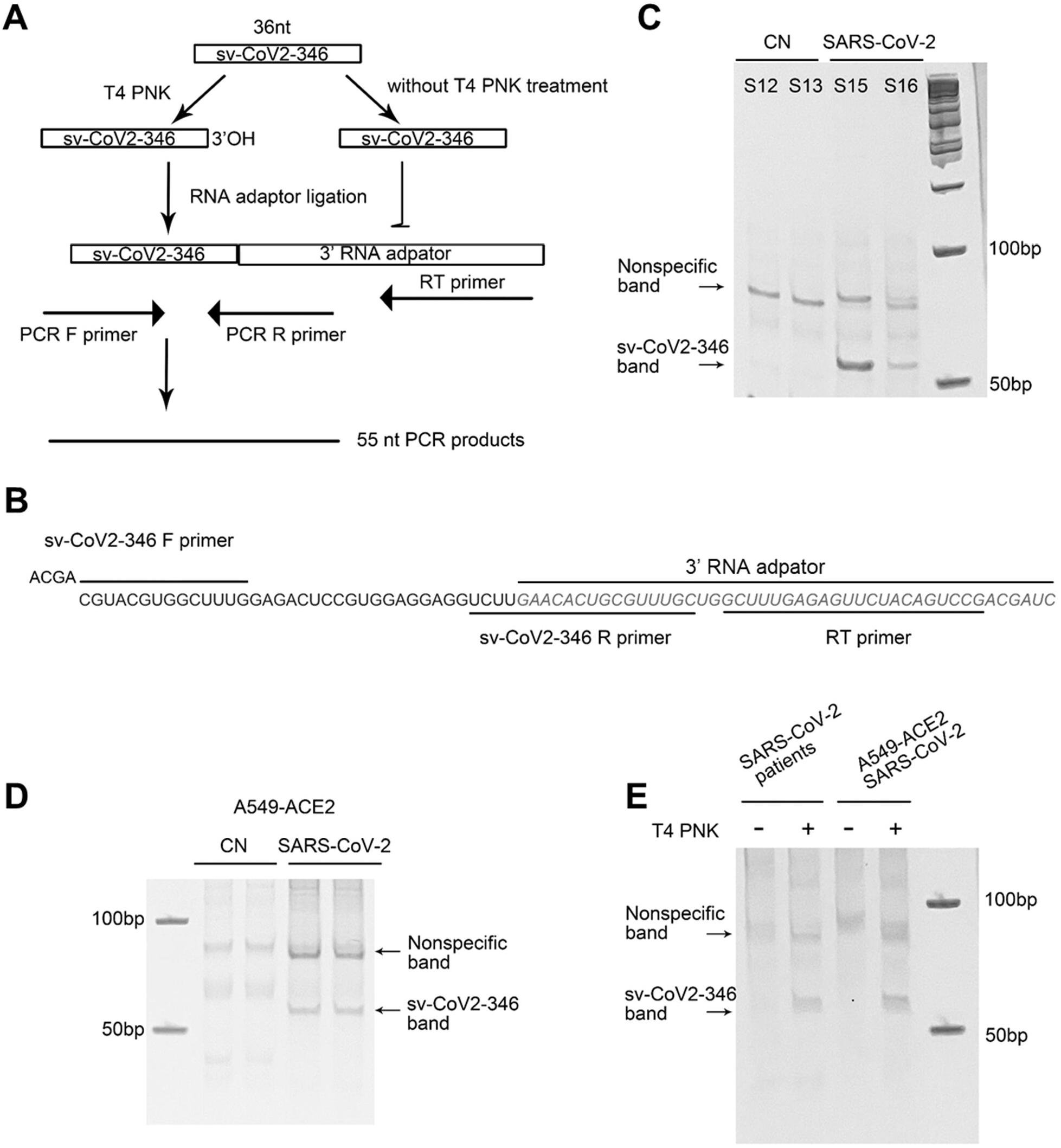
Experimental validation of sv-CoV2-346. **(A)** Schematic cartoon illustration on how to detect sv-CoV2-346 by RT-PCR and 3’-OH at the 3’-end. **(B)** The illustration of primers and the 3’ RNA linker for sv-CoV2-346 detection by RT-PCR. **(C)** RT-PCR was performed to detect sv-CoV2-346 in NPS samples of two CN and two SARS-CoV-2 patients. **(D)** Confirmation of sv-CoV2-346 in SARS-CoV-2-infected A549-ACE2 cells. **(E)** Detection of sv-CoV2-346 needs the pretreatment of T4 PNK. To 3’-end of sv-CoV2-346 is not –OH end, as samples without pretreatment of T4 PNK did not result in sv-CoV2-346 bands. All experiments were independently repeated twice.

Coronavirus-encoded svRNAs have been previously reported to be 18∼22 nt long and therefore, share similar lengths with miRNAs [28, 31]. Coronavirus-encoded svRNAs with lengths longer than 30 nt have not been identified. Herein, we think that the identification of additional SARS-CoV-2-derived svRNAs was benefited from the treatment of T4 PNK and RNA linker ligation at their 3’-end. As shown in **Figure 6E**, both patient or infected cell samples, without such treatments, did not result in the band presence, supporting the lack of 3’-OH end of sv-CoV2-346 and the necessity of specific T4 PNK treatment for sv-CoV2-346 dection.

Herein, we also initialed to characterize sv-CoV2-346 by predicting the secondary RNA structure of svRNAs. Besides sv-CoV2-346, there were two other svRNAs, sv-CoV2-299 and sv-CoV2-404, near the region where sv-CoV-346 was derived (**Figure. 7A**). sv-CoV2-299, sv-CoV2-346, and CoV2-404 were derived from nucleotide 299 to 328, 346 to 382, and 400 to 443, respectively. Therefore, we took the viral genome spanning from 289 to 485, which covers all these three regions with some nt extension on both ends and predicted its RNA secondary structure using RNAfold web server based on minimum free energy to have a clue of biogenesis mechanisms [48](**Figure. 7B**). We found that nucleotides 299, 328, 400, 443, and 382 are all located on loops, implying the cleavage at these five sites along with the single-stranded RNA by ribonuclease (**Figure. 7B**). Only nucleotide 346 was in the middle of the stem (**Figure. 7B**). Interestingly, we found that 68 nt long svRNAs (sv-CoV2-314) overlapped with sv-CoV2-346. We, therefore, took the genome section spanning from 314 to 382 and run the secondary and tertiary structures of sv-CoV2-314 using RNAfold web server and RNAComposer, respectively [48, 49](**Figures 7C and D**). This 68 nt fragment contained three hairpin loops and was folded into an L-shaped-like tertiary structure, and nucleotide 346 was located within the bottom loop (**Figures 7C and D**). The secondary and tertiary structures of sv-CoV2-314 were similar to tRNA, and nucleotide 346 location was similar to the cleavage site of tRFs (**Figures 7C-F**). Thus, we speculated that 68nt sv-CoV2-314 may be the precursor of 36nt sv-CoV2-346 and the virus may use endonuclease involved in tRF biogenesis to generate viral small RNAs fragments.

**Figure 7.**
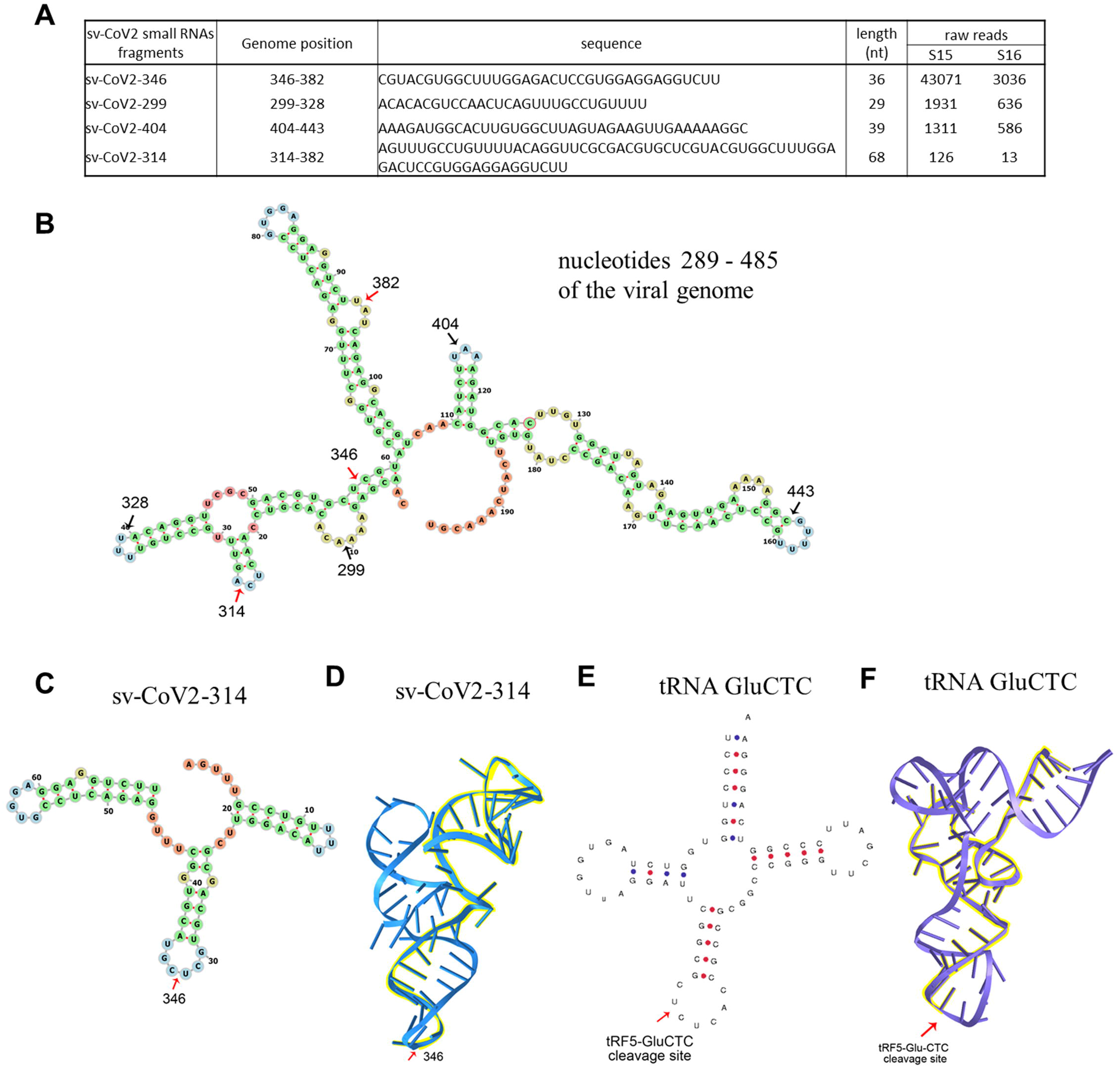
Structure prediction of viral small RNA. **(A)** The small RNA sequences with high raw reads mapping on viral nucleotides 289 – 485. **(B)** The secondary structure of viral nucleotides 289 – 485. The secondary **(C)** and tertiary **(D)** structure of 68nt sv-CoV2-314. The secondary **(E)** and tertiary **(F)** structure of a tRNA example-GluCTC. Arrows indicated cleavage sites.

## Discussion

In this study, we identified the change in sncRNA expression by SARS-CoV-2. sncRNAs include miRNAs, piRNAs, tRFs, and snoRNAs, etc. Among these, miRNAs have been getting lots of attention in the studies [50-53]. Standard barcode-ligation-based small RNA-seq are usually designed to capture miRNAs, which are sncRNAs cleaved from their precursors by RNase III class enzymes and usually present 3’-OH ends [54]. It is increasingly acknowledged that the 3’-ends of other types of sncRNAs are heterogeneous [32], resulting in unsuccessful sequencing barcode ligation and subsequently undetectable sncRNAs by standard small RNA-seq protocols. In our study, T4 PNK-RNA-seq was employed to profile sncRNAs with heterogeneous ends. Sequencing data revealed that piRNAs and tRFs had higher global expression than miRNAs in NPS (**Figure 1A**). It is possible that sncRNAs carry various unidentified modifications, which are insensitive to T4 PNK treatment, resulting in these sncRNAs undetectable still by T4 PNK-RNA-seq. Herein, the consistency among the seq data, qRT-PCR result, and NB data of tRF5-GluCTC suggested the reliability of T4 PNK-RNA-seq for tRF5 detection.

Notable, among differently expressed tRF5s with basemean > 20 in CN or SARS-CoV-2 groups (**Table V**), we found four tRF5s: tRF5-GluTTC-1-2, tRF5-GlyCCC-6-1, tRF5-LysCTT6-1 and tRF5-SecTCA-2-1, were not classic tRF5s. While their 3’-ends commonly stop around the anticodon region like classical tRF5s, they lack the first 10-15 nt of the tRNA 5’end. Since they span the complete region of the <u>D</u> loop and the first half of the anti<u>c</u>odon loop, we subgrouped and named them as tRF5DC (**Supplementary Figure 1**). Interestingly, tRF5DC and classic tRF5s were derived from the different tRNA isoacceptors tRNA, suggesting different biogenesis mechanisms of these two type tRFs.

This study further supported that tRF induction is virus-specific. Previously, we and others have shown that RSV, hepatitis B virus (HBV), and hepatitis C virus (HCV) infections lead to different tRF profile changes [21, 22, 37]. Compared with tRF induction by RSV, we found that SARS-CoV-2 could induce tRF5-ValCAC, while RSV cannot. On the other hand, we found that tRF5-GlyGCC, which is significantly inducible by RSV, was not induced by SARS-CoV-2.

Other than host-derived sncRNAs, sncRNAs can also be derived from viruses. svRNAs have been reported to be involved in the regulation of viral replication, viral persistence, host immune evasion, and cellular transformation [27]. SARS-CoV-2-encoded sncRNAs have been vitrificated by two independent groups [31, 55]. Zheng F et.al found SARS-CoV-2-derived miR-nsp3-3p served as a serum biomarker to prioritize the patients at high risk of developing the severe disease [55]. Our sequencing data revealed remarkable svRNAs reads in NPS samples of SARS-CoV-2 patients and the abundance of these svRNAs varied among individuals. sv-CoV2-346 was also present in SARS-CoV-2 infected A549-ACE2 cells. Due to the limited NPS RNA samples, the leftover RNAs, after sequencing and qRT-PCR validation for tRF5s, were not enough for stem-loop qRT-PCR for svRNAs. Thus, we detected sv-CoV2-346 by RT-PCR using the same cDNA generated by the RT step for the qRT-PCR assays for tRF5s. Our RT-PCR, revealed a sv-CoV2-346 specific band around 55 nt and a non-specific band. In the future, we will study the relationship between SARS-CoV-2 svRNAs and viral loads/disease severity.

The most widely studied viral sncRNAs are miRNAs-like svRNAs. These ∼20nt viral miRNAs can function as miRNAs to regulate host and viral gene expression. Both DNA and RNA viruses could encode miRNAs-like svRNAs via Dicer-dependent miRNAs biogenesis pathways [27, 56]. Among RNA viruses, cytoplasmic restricted RNA viruses were thought incapable of producing miRNA-like svRNAs. However, accumulating evidence indicates cytoplasmic RNA viruses, such as H5N1 influenza virus, enterovirus 71(EV71), West Nile virus (WNV), SARS-CoV, and SARS-CoV-2, also encode viral miRNAs [28, 31, 55, 57-60]. These cytoplasmic RNA viruses generate viral miRNAs via multiple non-canonical miRNAs biogenesis mechanisms. Dicer, not Drosha, is involved in WNV and EV71 viral miRNAs generation [59, 60]. H5N1 influenza virus and SARS-CoV encode viral miRNAs also in a Dicer-and Drosha-independent way [28, 58]. Besides viral miRNAs, the induction of functional svRNAs which do not look like miRNAs was reported for cytoplasmic RNA viruses [57]. Overall, the knowledge on cytoplasmic RNA viruses-encoded sncRNAs is limited. In terms of SARS-CoV-2-derived svRNAs, several SARS-CoV-2 svRNAs were reported [31, 55]. In this study, we revealed new SRAS-CoV-2 svRNAs, all of which were longer than miRNAs. sv-CoV2-346, the most abundant svRNAs, seemed to lack the 3’-OH, as the detection needs T4 PNK treatment. Moreover, we found 68nt sv-CoV2-314 may have a similar tertiary structure as tRNAs and may function as the potential precursor of 36nt sv-CoV2-346 (**Fig. 7**). How viruses use the host proteins to generate viral small RNAs fragments is still unclear and awaits investigation.

In summary, we investigated COVID-19-impacted sncRNAs comprehensively using the NPS samples by T4 PNK-RNA-seq and modified qRT-PCR method. We are aware that our study has some limitations, such as T4 PNK-RNA-seq cannot restore and catch all the type ncRNAs. Although the sample size of patients is limited, our methods enabled us to identify tRF signatures of SARS-CoV-2 infection. Our methods also enabled us to find several new SARS-CoV-2-encoded sncRNAs, whose functions and biogenesis will be studied in the near future.

## Legend

**Supplementary Figure 1**. Biogenesis of classic tRF5s and tRF5DC.

## Author Contributions

W.W. and X.B. wrote the manuscript; W.W., EJ. C., B.W., K Z., contributed to the experiment performance and data analyses. L.T., P. H., R. G., I. I., L. H., and I. L contribute to the deep seq data analyses and related methodology establishment. G.H. and J.D. developed IBM protocols for sample collection and NSP sample handling. J.D., T.W., and X.B. contributed to overall project management and experimental design.

## Acknowledgments

We thank Mrs. Darby L. Buck for the manuscript editing.

