## Supplementary figures and images for "Changes of small non-coding RNAs by severe acute respiratory syndrome coronavirus 2 infection"

### Supplemental Figure 1

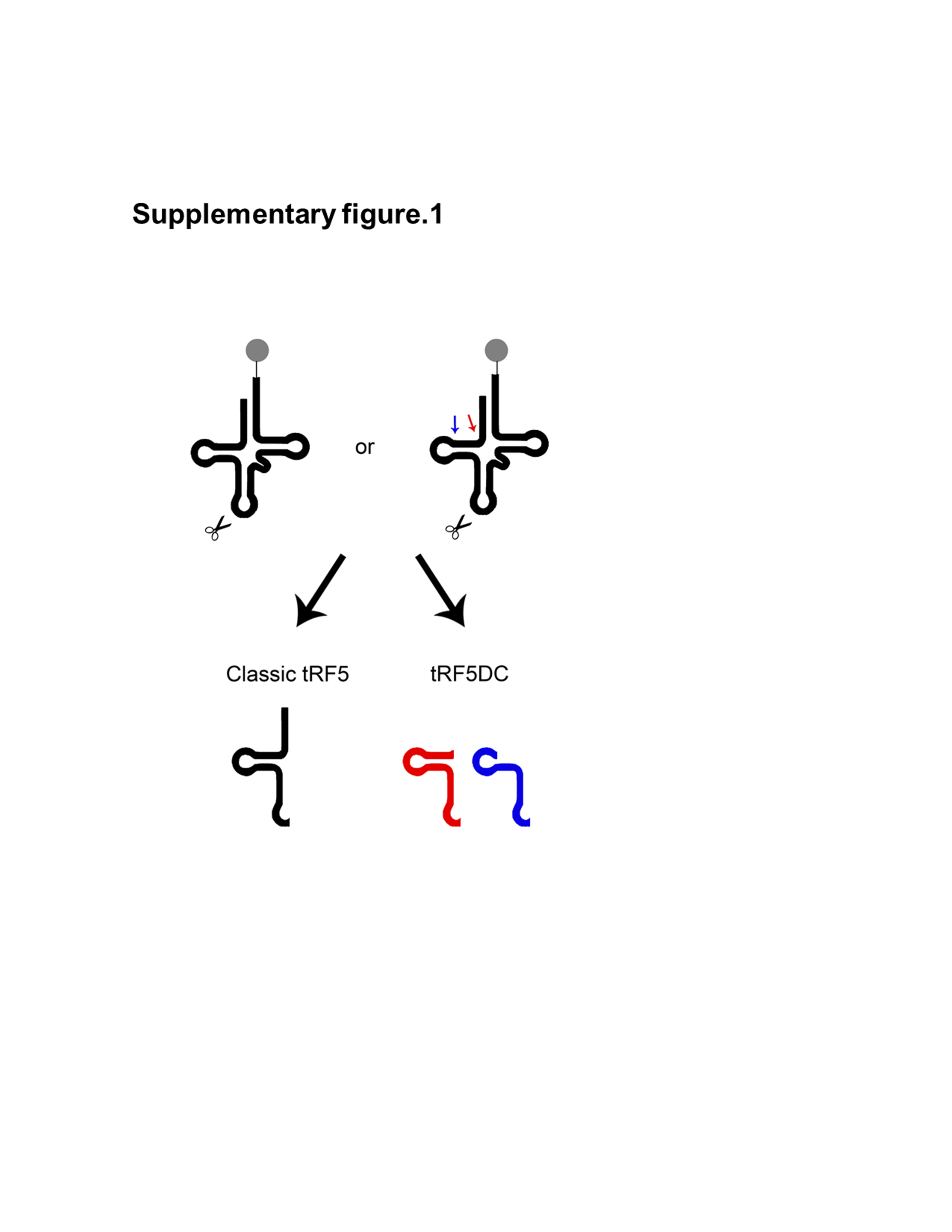
