## Supplemental Table 1 for "Changes of small non-coding RNAs by severe acute respiratory syndrome coronavirus 2 infection"

| | BaseMean |
|---|---|
| piR-31038 | 13119.37 |
| piR-38581 | 5149.83 |
| piR-38756 | 3153.08 |
| piR-55891 | 2726.51 |
| piR-33382 | 2348.61 |
| piR-59293 | 2231.93 |
| piR-59425 | 1488.54 |
| piR-58707 | 1416.84 |
| piR-51761 | 1292.06 |
| piR-33043 | 1249.65 |

**Supplementary Table I**. The Top 10 piRNAs in NPS samples from SARS-CoV-2 patients
