## Supplemental Table 2 for "Changes of small non-coding RNAs by severe acute respiratory syndrome coronavirus 2 infection"

| | baseMean |
|---|---|
| tRF5-Glu-CTC-2-1 | 45637.69747 |
| tRF5-Gly-GCC-3-1 | 4785.065821 |
| tRF5-Glu-TTC-8-1 | 888.8149209 |
| tRF5-Gly-GCC-1-5 | 750.6580236 |
| tRF5-Val-CAC-chr1-93 | 467.6325707 |
| tRF5-nm-Tyr-GTA-chr14-8 | 398.5972534 |
| tRF5-Lys-CTT-2-5 | 354.265456 |
| tRF5-SeC-TCA-2-1 | 325.3529339 |
| tRF5-Glu-TTC-chr1-138 | 270.7106895 |
| tRF5-His-GTG-1-8 | 194.8375191 |

**Supplementary Table II.** The Top 10 tRFs in NPS samples

| sequence |
| --- |
| UCCCUGGUGGUCUAGUGGUUAGGAUUCGGCGCU |
| GCAUUGGUGGUUCAGUGGUAGAAUUCUCGCC |
| UCCCCUGUGGUCUAGUGGUUAGGAUUCGGCGCU |
| GCAUGGGUGGUUCAGUGGUAGAAUUCUCGCC |
| GUUUCCGUAGUGUAGUGGUUAUCACGUUCGCU |
| GCUGAGUGAAGCAUUGGACUGUAA |
| GCCCGGCUAGCUCAGUCGGUAGAGCAUGAGACU |
| AGUGGUCUGGGGUGC |
| UCCCUGGUGGUCUAGUGGCUAGGAUUCGGCGCU |
| GGCCGUGAUCGUAUAGUGGUUAGUACUCUGCGUU |

; from SARS-CoV-2 patients
